# Imprinting effects of UBE3A loss on synaptic gene networks and Wnt signaling pathways

**DOI:** 10.1101/649491

**Authors:** S. Jesse Lopez, Benjamin I. Laufer, Ulrika Beitnere, Elizabeth L. Berg, Jill L. Silverman, David J. Segal, Janine M. LaSalle

## Abstract

The genomically imprinted *UBE3A* gene encodes a E3 ubiquitin ligase whose loss from the maternal allele leads to the neurodevelopmental disorder Angelman syndrome. However, the mechanisms by which loss of maternal UBE3A contribute to severe neurodevelopmental phenotypes are poorly understood. Previous studies of UBE3A function have focused on mouse models or single targets, but these approaches do not accurately reflect the complexity of imprinted gene networks in the brain nor the systems-level cognitive dysfunctions in Angelman syndrome. We therefore utilized a systems biology approach to better elucidate how UBE3A loss impacts the early postnatal brain in a novel CRISPR/Cas9 engineered rat Angelman model of a complete *Ube3a* deletion. Strand-specific transcriptome analysis of offspring derived from maternally or paternally inherited *Ube3a* deletions revealed the expected parental expression patterns of *Ube3a* sense and antisense transcripts by postnatal day 2 (P2) in hypothalamus and day 9 (P9) in cortex, when compared to wild-type sex-matched littermates. The dependency of genome-wide effects on parent-of-origin, *Ube3a* genotype, and time (P2, P9) was investigated through transcriptome (RNA-seq of cortex and hypothalamus) and methylome (whole genome bisulfite sequencing of hypothalamus). Weighted gene co-expression and co-methylation network analyses identified co-regulated networks in maternally inherited *Ube3a* deletion offspring correlated with postnatal age that were enriched in developmental processes including Wnt signaling, synaptic regulation, neuronal and glial functions, epigenetic regulation, ubiquitin, circadian entrainment, and splicing. Furthermore, using this novel rat model, we showed that loss of the paternally expressed *Ube3a* antisense transcript resulted in both unique and overlapping dysregulated gene pathways, predominantly at the level of differential methylation, when compared to loss of maternal *Ube3a*. Together, these results provide the most holistic examination to date of the molecular impacts of UBE3A loss in brain, supporting the existence of interactive epigenetic networks between maternal and paternal transcripts at the *Ube3a* locus.

**Author Summary:** The neurodevelopmental disorder Angelman syndrome is caused by loss of *UBE3A* from the maternal chromosome. *UBE3A* is a genomically imprinted gene, which results in parent-of-origin specific expression of a protein from the mother and a noncoding RNA from the father. While mouse models have been useful in investigating diverse roles for UBE3A, their partial mutations are of limited utility for investigating parental imprinting effects or identifying a complete list of downstream differences in gene pathways relevant to developing therapies for Angelman syndrome. To address this limitation, we utilized a novel rat model with a CRISPR/Cas9 engineered full UBE3A deletion and systems biology approaches to better understand how UBE3A loss affects early postnatal brain development. We discovered that UBE3A loss has widespread effects on many important neuronal and cellular pathways and uncovered interesting interactions between maternal and paternal genes that were not previously considered. Taken together, our findings provide the most comprehensive view of UBE3A’s influences in the brain, which are relevant to the understanding and development of treatments for Angelman syndrome and related neurodevelopmental disorders.

## Introduction

*UBE3A* encodes an E3 ubiquitin ligase that targets multiple proteins for proteasomal degradation (1,2). Residing within the human 15q11.2-q13.3 locus, *UBE3A* is parentally imprinted, specifically in mature neurons but not glia, where it becomes silenced on the paternal allele due to the paternal-specific expression of an anti-sense transcript (*UBE3A-ATS*) originating from an unmethylated imprinting control region on the paternal allele. Angelman syndrome (AS) is a neurogenetic disorder caused by deletion of the maternal 15q11.2-q13.3 locus or maternal-specific *UBE3A* mutation (3). However, a large population-based study recently demonstrated that both maternal and paternal duplications of 15q11.2-q13.3 are associated with increased risk of autism spectrum disorders (ASD) or developmental delays (4). Paternal transcripts expressed within human 15q11.2-q13.3 include a long polycistronic transcript encoding the splicing protein SNRPN, snoRNA host genes with independent functions that also produce multiple non-coding snoRNA subunits, as well as *UBE3A-ATS*. Loss of the paternally expressed *SNORD116* locus causes Prader-Willi syndrome and results in circadian dependent changes in gene expression and DNA methylation genome-wide in cortex (5,6), as well as cognitive impairments (7). However, the mechanisms behind how the similarly regulated paternal *UBE3A-ATS* transcript may contribute to endophenotypes is currently unknown.

Recent research has illuminated diverse roles for UBE3A in neuronal and cellular functions. Proteomic analyses have suggested that UBE3A and its interacting proteins integrate several cellular processes including translation, intracellular trafficking, and cytoskeleton regulation, which are all necessary for neuronal function (8,9). In line with its role in neurodevelopmental disease, UBE3A has also been shown to be critical in synaptic formation and maintenance (10–12). We have also recently examined the nuclear chromatin-related genome-wide effects of *UBE3A* dysregulation in neurons, which revealed significant effects on DNA methylation profiles that were characterized by differentially methylated regions (DMRs) in genes involved in transcriptional regulation and brain development (13). How these diverse functions might contribute to *UBE3A*-related disorders *in vivo* has yet to be explored.

Over the past decade, rapid progress has been made in understanding biological networks formed by complex sets of interactions between numerous coding and non-coding genes that play important role in deciphering disease phenotypes (14,15). Gene co-expression analysis in particular was designed to identify correlated patterns of gene expression across different experiments, tissues, or species (16–19). In particular, gene expression networks have been increasingly used to obtain systematic views about an immensely complex molecular landscape across brain development (20–25). One recent approach for analyzing gene co-expression networks is to identify Topological Overlap between functional modules or subnetworks that are relevant to the disease. Module discovery using the Weighted Gene Co-expression Network Analysis (WGCNA) package (17,26) is a widely utilized method for this purpose. WGCNA aims to identify modules of genes that are highly correlated based on their co-expression patterns and does not depend on statistically significant differential expression at the single gene level. The usefulness of this systems biology method has been widely demonstrated as evidenced by multiple publications from a broad range of diseases such as cancer, epilepsy, neurodegenerative and neurodevelopmental disorder (25,27–29).

To gain stronger insight into how the diverse functions of *UBE3A* influence neurodevelopment, we leveraged a systems biology approach to examine a novel *UBE3A* CRISPR-deletion AS rat model and integrated the transcriptome (RNA-seq) and methylome (WGBS). By identifying the genome-wide changes in the developing brain caused by deletion of *Ube3a*, we uncovered networks and pathways affected by *UBE3A* at both the transcriptomic and epigenomic levels, including neuronal and synaptic processes and critical cellular pathways. In addition, while the overwhelming majority of reports focus solely on the maternal deletion, we utilized reciprocal breeding, so that both the maternal and paternal-derived deletions were investigated, revealing genome-wide influences of both maternal *Ube3a* and paternal *Ube3a-Ats* transcripts. Adding to the scope, we examined two brain regions at two developmental time points.

## Results

### The novel CRISPR/Cas9 knockout rat model contains a complete *Ube3a* deletion that enables allele-specific investigations

While *Ube3a* mutant mice have been extensively used as models of AS (30–34), none of these are complete knockouts of the gene. To produce a more robust model of AS and to better understand the functional consequences of maternal UBE3A loss, an AS rat model was designed to completely delete *Ube3a* by targeting CRISPR/Cas9 nucleases to sites flanking *Ube3a* (S1 Figure).

We devised a breeding strategy to produce both maternally and paternally-derived *Ube3a* deletions and harvested hypothalamus and cortex from deletion animals and litter-matched controls at postnatal day 2 (P2) and postnatal day 9 (P9) (**Fig 1**). These two timepoints were chosen because *Ube3a* was expected to become imprinted within the first week of life, based on prior studies in mouse (35,36), and this was also the time window when deficits in pup ultrasonic vocalization and developmental delays were observed in maternal *Ube3a* deletion rat pups when compared to wild-type littermates (37). RNA-seq browser tracks for hypothalamus (**Fig 2A**) and cortex (**Fig 2B**) confirmed the expected loss of the *Ube3a* sense transcript (blue) in maternal *Ube3a* deletion (Mat−/+) samples at both time points when compared to Mat+/+ wild-type. In contrast, paternally inherited *Ube3a* deletion (Pat+/-) hypothalamus and cortex samples exhibited the complete loss of the *Ube3a-Ats* transcript (red) when compared to wildtype. In maternally-derived *Ube3a* deletion hypothalamic P2 and P9 samples, expression from the *Ube3a* exons is reduced to ∼10% of wild-type control, which is consistent with the expected biallelic expression in glia. In contrast, expression of *Ube3a-Ats* over the *Ube3a* gene body is completely lost in the paternally-derived deletion samples, which was expected based on the imprinted expression in all tissues. In the hypothalamus, expression of *Ube3a-Ats* was equivalent at both the P2 and P9 time points, indicating that *Ube3a* is imprinted prior to P2 in rat hypothalamus, consistent with recent estimates in mouse (38). In P2 cortex, however, imprinted expression of *Ube3a* was less complete than in P9 cortex or P2 hypothalamus, which is also consistent with findings in mouse of a delayed onset of developmental *Ube3a* imprinting in cortex compared with sub-cortical brain regions.

**Fig 1.**
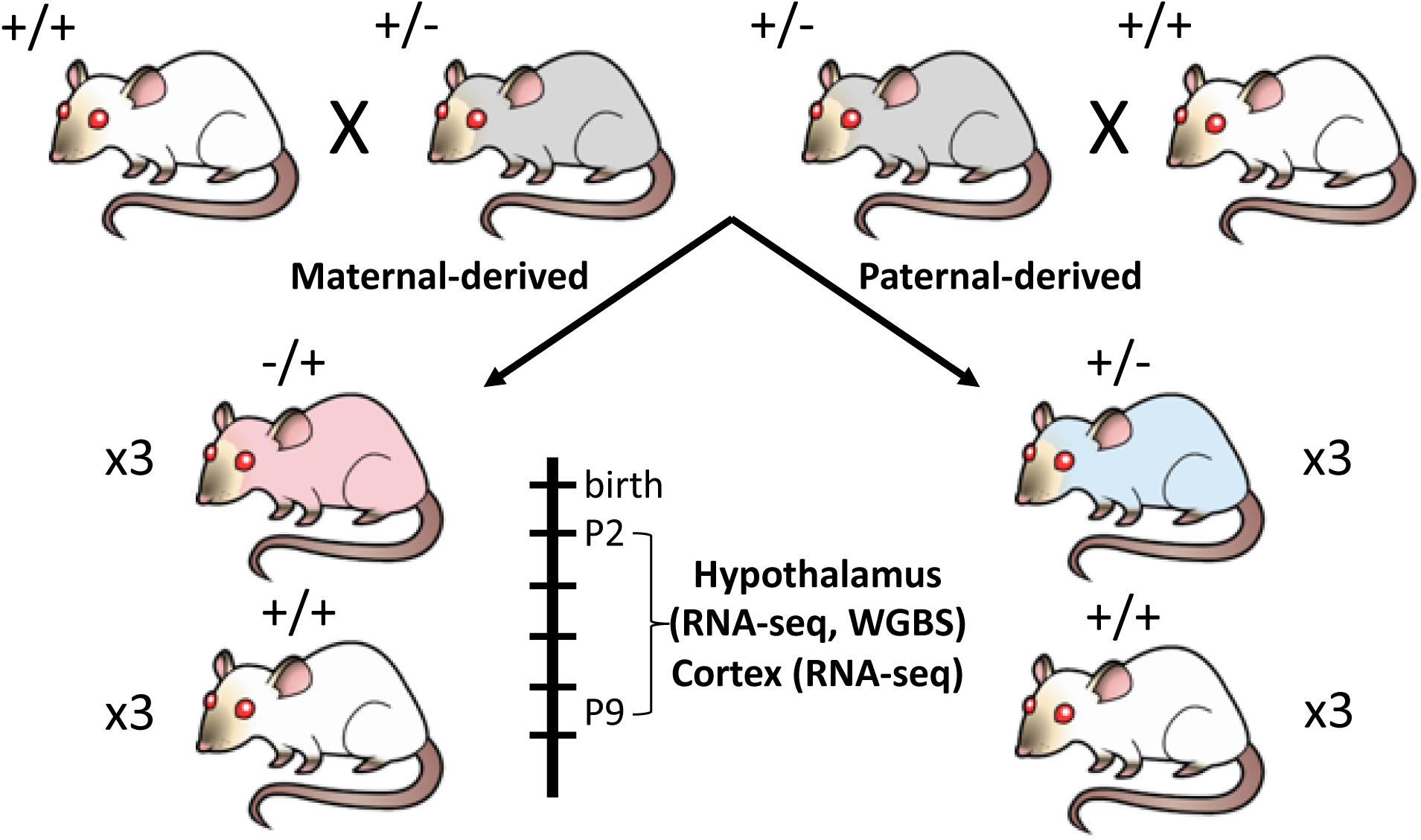
*Ube3a* CRISPR-deletion rat model study design. Reciprocal crosses with *Ube3a*-deletion carriers (grey) produced the maternal-derived (red) and paternal-derived (blue) deletion. Hypothalamus and cortex were harvested from female rat pups at P2 and P9 in triplicate with litter-matched controls and utilized for RNA-seq and WGBS.

**Fig. 2.**
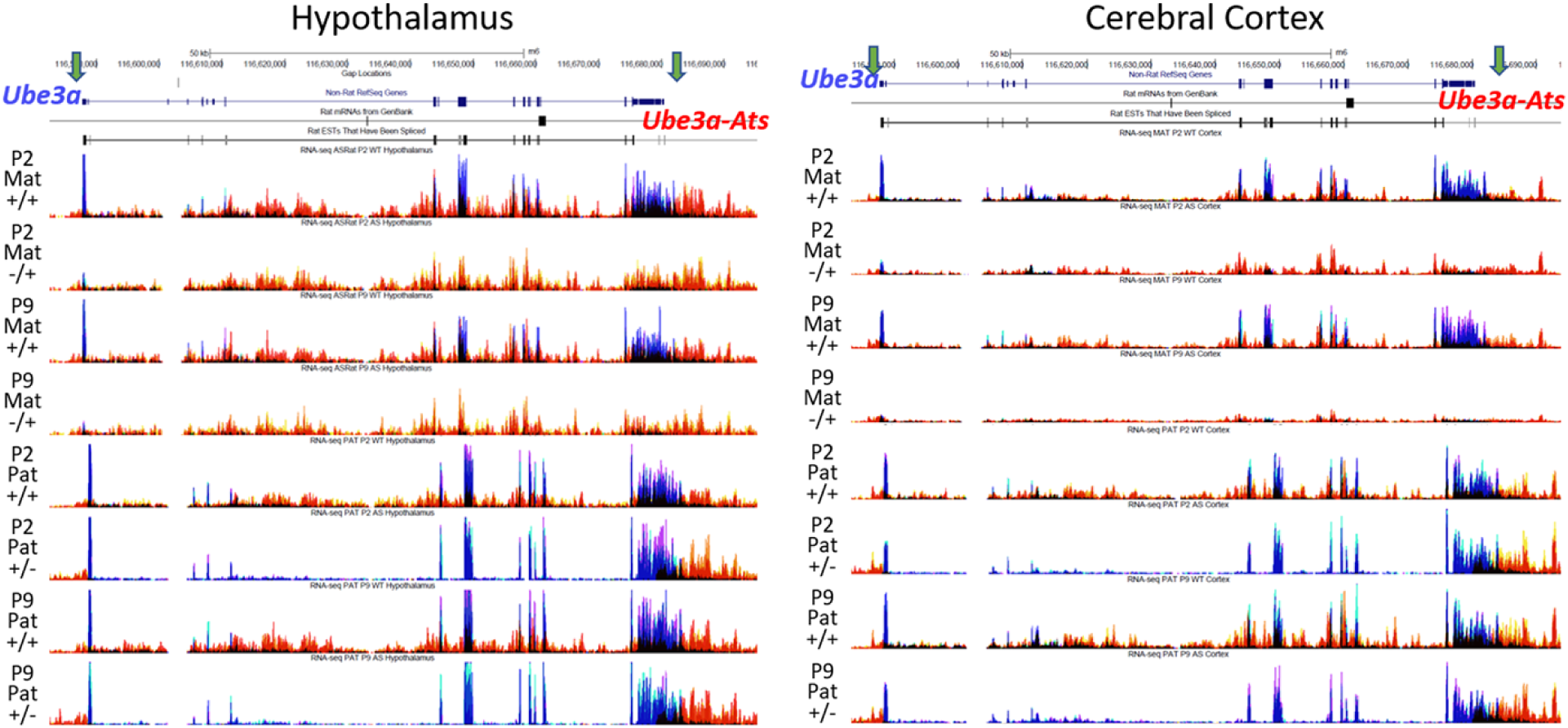
Imprinting patterns of *Ube3a* and *Ube3a-Ats* observed by strand-specific RNA-seq tracks. RNA-seq tracks at the *Ube3a* locus for hypothalamus (left) and cortex (right) samples showing sense-strand (blue) and anti-sense-strand (red) expression. Maternal deletion samples (P2 Mat−/+ and P9 Mat−/+) displayed a loss of exonic *Ube3a* whereas paternal deletion samples (P2 Pat−/+ and P9 Pat−/+) showed a loss of *Ube3a-Ats* over the deleted *Ube3a* gene body.

### Identification of dysregulated gene functions and pathways by *Ube3a* genotype, parental origin, and postnatal age from RNA-seq and WGBS

To identify genes dysregulated at the transcript level as a result of *Ube3a* deletion across developmental time, we first performed a standard pair-wise differential gene expression (DGE) analysis using four comparison groups based on time point or parental genotype (S2 Table, S3 Figure, S4 Data). While *Ube3a* and a few other transcripts reached genome-wide significance for differential expression based on genotype, the complexities in our experimental design to examine genotype parent-of-origin and time interactions were overall not well served by this standard pair-wise bioinformatic approach. Similarly, we performed pair-wise differentially methylated region (DMR) analyses for either genotype or time using the other as a covariate (S5 Table, S6 Figure, S7 Figure, S8 Data), revealing a similarly large number of pair-wise DMRs across the entire genome.

Because of the likelihood for interactions of parent-of-origin imprinting effects and postnatal age with *Ube3a* deletion, we sought to improve the analysis of transcriptomic and methylomic data using a systems biology approach. We therefore utilized WGCNA to first identify groups of co-expressed or co-methylated genes and then measure their correlation with either postnatal age or *Ube3a* genotype (sample cluster dendrograms in S9 Figure). We used a litter-based approach where we performed independent WGCNAs on data from either the maternal or paternal derived litters. In addition to wild-type littermate controls being included in litter-based WGCNAs, control WGCNAs were also performed on wild-type offspring across maternal versus paternal derived litters as a comparison for background litter effects.

Co-expressed gene modules were detected and tested for correlation with either time point or genotype in separate analyses of hypothalamus (**Fig 3**) and cortex (S10 Figure). Positively-correlated modules in red represent increased expression or methylation in *Ube3a* deletion samples compared to wild-type or P9 samples compared to P2. Correlations colored in blue were negative, indicating decreased expression or methylation in *Ube3a* deletion samples or at P9 relative to P2. We then focused on the modules with *p*<0.05, shown by yellow frames. A summary of the number of WGCNA modules identified and the number passing this significance cutoff is shown in **Table 1**.

**Table 1.** WGCNA module summary

| Tissue | Deletion parent of origin | Assay | Input genes/ regions | Total number of modules | Significant genotype modules (neg:pos) | Genotype genes (neg:pos) | Significant time modules (neg:pos) | Time genes (neg:pos) |
| --- | --- | --- | --- | --- | --- | --- | --- | --- |
| Hypoth | Maternal | RNA | 25562 | 26 | 1:1 | 149:294 | 1:2 | 672:1006 |
|  |  | Methyl | 18335 | 45 | 4:4 | 1587:2028 | 4:4 | 1309:1858 |
|  | Paternal | RNA | 34812 | 18 | 0:0 | 0:0 | 2:1 | 2354:194 |
|  |  | Methyl | 13874 | 32 | 2:4 | 726:1534 | 4:2 | 2011:875 |
|  | Control | RNA | 27730 | 18 | 0:3 | 0:2774 | 0:3 | 0:1361 |
|  |  | Methyl | 16243 | 0 | - | - | - | - |
| Cortex | Maternal | RNA | 34470 | 18 | 1:0 | 1209:0 | 5:1 | 4301:5389 |
|  | Paternal | RNA | 33273 | 38 | 1:1 | 150:156 | 4:3 | 4725:1009 |
|  | Control | RNA | 34318 | 20 | 1:0 | 2271:0 | 0:0 | 0:0 |

**Fig 3.**
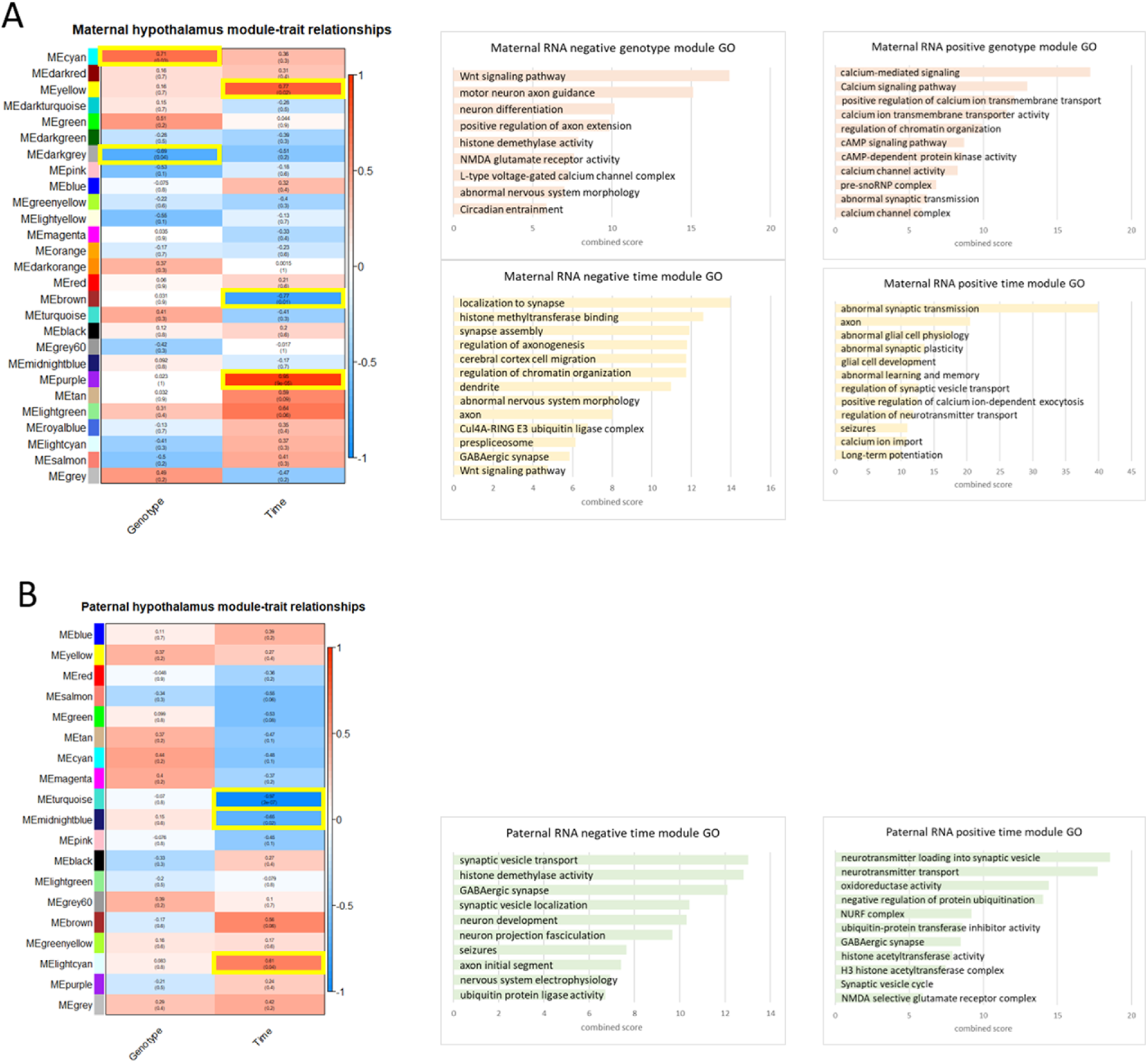
WGCNA expression module and trait correlation heatmaps and GO terms. Co-expression modules detected from network analysis of RPKM in **A)** maternal and **B)** paternal deletion hypothalamus. Modules were grouped by co-expression and tested for correlation with the traits of genotype or time. Modules of interest are correlated (red) or anticorrelated (blue) with genotype or time (p-value < 0.05, highlighted yellow boxes).Highlighted co-expression module gene lists were used for GO enrichment with Enrichr using GO Biological Process 2018, GO Cellular Component 2018, GO Molecular Function 2018, KEGG 2016, and MGI Mammalian Phenotype Level 4 categories. Top GO categories are displayed as combined score, calculated as -log10(p-value)*z-score.

To gain insight into the pathways and networks affected by genotype and time respectively, these trait-associated modules were used for gene ontology (GO) enrichment. For negatively-correlated modules, samples obtained from maternal *Ube3a* deletion and wild-type littermates contained 1 module of 672 co-expressed genes with decreased expression from P2 to P9 in the hypothalamus. Genes within this “maternal negative time module” were enriched in synaptic regulation, epigenetic regulation, axonogenesis, neuron maintenance, ubiquitin activity, splicing, and Wnt signaling (**Fig 3A**). Two positively-correlated maternal time modules containing 1006 genes were enriched in genes with functions in synaptic plasticity and synaptic transmission, glial development, and learning and memory. The paternally derived *Ube3a* deletion litters yielded two negatively-correlated time modules with 2354 genes in hypothalamus and one positively-correlated module of 194 genes that were similarly enriched in synaptic regulation, epigenetic regulation, neuron development, and ubiquitin activity (**Fig 3B**). As expected, significant genotype modules were only detected in maternal samples from WGCNA and were enriched in Wnt signaling, neuron differentiation and maintenance, epigenetic regulation, synaptic activity, and circadian entrainment for negatively-correlated modules, and calcium signaling, chromatin organization, and cAMP signaling for positively correlated modules (**Fig 3A**). GO enrichment for negatively-correlated modules detected in cortex samples were similar to those observed in hypothalamus modules for time, but the maternal genotype module was distinct in highlighting ubiquitin functions, dendrite, and spliceosome in cortex (S10 Figure). A control data set comparing wild-type samples from maternal and paternal litters produced three significant modules for parent-of-origin in hypothalamus and one in cortex with genes enriched in ubiquitin activity (S11 Figure). GO enrichment terms identified by WGCNA were highly similar to those observed by the simpler pairwise analysis (S12 Figure).

For WGCNA of methylation in the hypothalamus, we used the testable “background” regions of potential significant differential methylation from dmrseq to identify gene modules associated with differential methylation by genotype or time (**Fig. 4**). Co-methylated gene modules were identified similarly to the co-expressed gene modules, focusing on gene modules with the highest confidence correlations for determining GO term enrichments. In contrast to the co-expression analyses in which significant genotype effects were limited to maternal inheritance, multiple co-methylated gene modules based on genotype were observed in both maternal and paternal *Ube3a* deletion litters.

**Figure 4.**
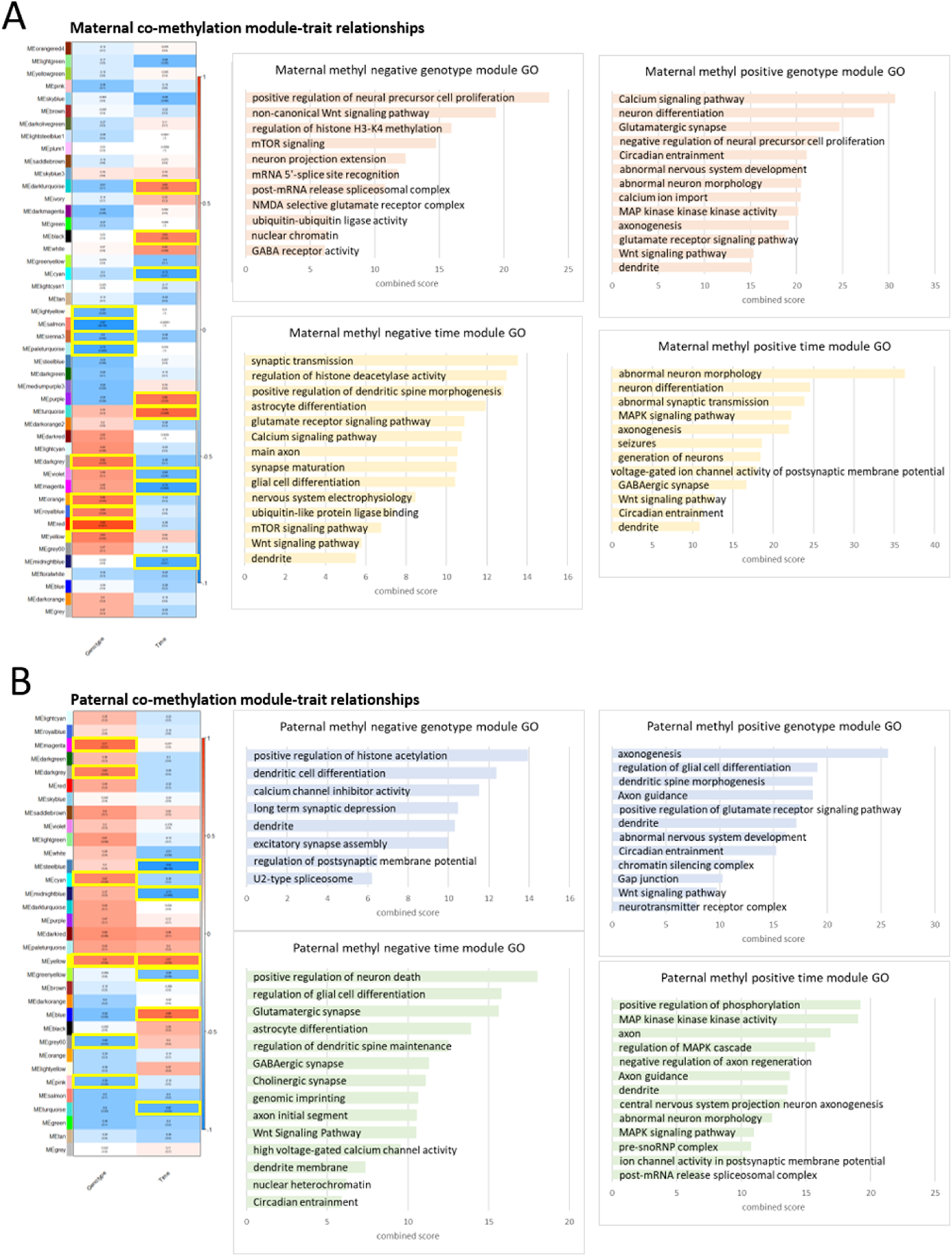
WGCNA methylation module and trait correlation heatmaps and GO terms. Co-methylation modules detected from background smoothed methylation values in maternal and paternal deletion hypothalamus. Modules were grouped by co-expression and tested for correlation with traits of genotype or time. Modules of interest are correlated (red) or anticorrelated (blue) with each trait (p-value < 0.05, highlighted yellow boxes). Highlighted co-expression module gene lists were used to determine GO enrichment with Enrichr using GO Biological Process 2018, GO Cellular Component 2018, GO Molecular Function 2018, KEGG 2016, and MGI Mammalian Phenotype Level 4 categories. Top GO categories are displayed as combined score, calculated as -log10(p-value)*z-score. Significant module gene lists can be found in S14 Data.

GO analysis of associated genes from co-methylated modules revealed distinctions in gene functions based on maternal versus paternal genotype effects. There were four genotype modules with 1366 co-methylated regions detected in maternal samples, which are enriched in neural precursor proliferation, non-canonical Wnt signaling, histone methylation, mTOR, splicing, synaptic activity, and ubiquitin activity (**Fig 4A**). Whereas, two genotype modules with 748 co-methylated regions in paternal samples were identified and were enriched in histone acetylation, excitatory synapse regulation, and splicing (**Fig 4B**). Time-associated modules were enriched in excitatory synaptic regulation, histone deacetylation, glial differentiation, ubiquitin activity, mTOR, and Wnt signaling for maternal samples, while paternal time modules were enriched for similar functions but also included imprinting and circadian entrainment. In contrast, the control co-methylation WGCNA from wildtype maternal versus paternal samples not only produced no significant modules, but no clustered modules were detected. Furthermore, GO enrichment terms identified by WGCNA were highly similar to those observed by the pair-wise analysis of differentially methylated genes (S13 Figure).

To directly compare the WGCNA results for the co-expression modules and co-methylation modules as well as across parent-of-origin of *Ube3a* deletion, we performed cluster profiling of the gene ontology results for each analysis. For individual GO terms, four distinct clusters were observed for downregulated (negatively-correlated modules) specific GO terms (S15 Figure), while the upregulated (positively-correlated modules) GO profile clustered into two groups (S16 Figure). Cluster profiling by KEGG pathways revealed five distinct clusters for downregulated pathways (**Fig 5A**): Group I clusters by downregulated pathway terms that were specific to maternal, but not paternal, genotype effects, some of which also overlapped with maternal genotype and time-associated co-methylation and paternal time-associated co-methylation terms. These included Wnt signaling, axon guidance, and other developmental processes and cancer. Group II clusters by downregulated pathways specific to maternal time-associated co-expression, aside from some associated with paternal time effects. These predominantly included longevity traits as well as insulin, ubiquitin, choline pathways, which also include genes in RNA metabolic processes. Group III downregulated pathways were uniquely associated with paternal time-associated co-expression, some of which also overlapped with time-associated co-methylation terms. These included GABAergic synapse and genes associated with morphine addiction. Groups IV and V are defined by downregulated pathways uniquely identified by co-methylation, but not co-expression WGCNAs. Groups IV and V are distinct in KEGG pathways either largely overlapping by maternal genotype and time (Group IV) or only time-associated but indiscriminate to parental origin (Group V). Group IV included major signal transduction pathways such as MAPK, cAMP, PI3K-Akt, and cGMP-PKG, while Group V included pathways related to functions in circadian entrainment, including insulin and oxytocin secretion. KEGG cluster profile for upregulated (positively-correlated) pathways grouped into four patterns (**Fig 5B**): Group I clusters the pathways upregulated by maternal *Ube3a* deletion genotype that also overlap with all co-methylation data sets. These pathways are enriched in signaling including calcium, adrenergic and cAMP signaling, as well as serotonergic synapse. Group II clusters the pathways upregulated with time and maternal inheritance that include long-term potentiation, as well as apelin signaling and gastric acid secretion. Groups III and IV clusters pathways commonly upregulated across the co-methylation analyses. Group IV upregulated pathways do not overlap with those identified by maternal genotype and include genes involved in GABAergic and cholinergic synaptic transmission in addition to signaling pathways such as FoxO, Ras, and Rap1. Group III clusters the most diverse of the upregulated pathways, which are enriched for the glutamatergic synapse, MAPK signaling, circadian entrainment, Wnt signaling, pluripotency, axon guidance, and oxytocin signaling. Paternal time co-expression pathways did not significantly overlap upregulated pathways but also included GABAergic synapse and synaptic vesicle cycle (S17 Data). This pathway clustering analysis supports the hypothesis that maternal *Ube3a* genotype effects are predominantly regulating transcriptional differences over time in this AS rat model, while dysregulation in DNA methylation involves contributions of both maternal and paternal *Ube3a* alleles.

**Fig 5.**
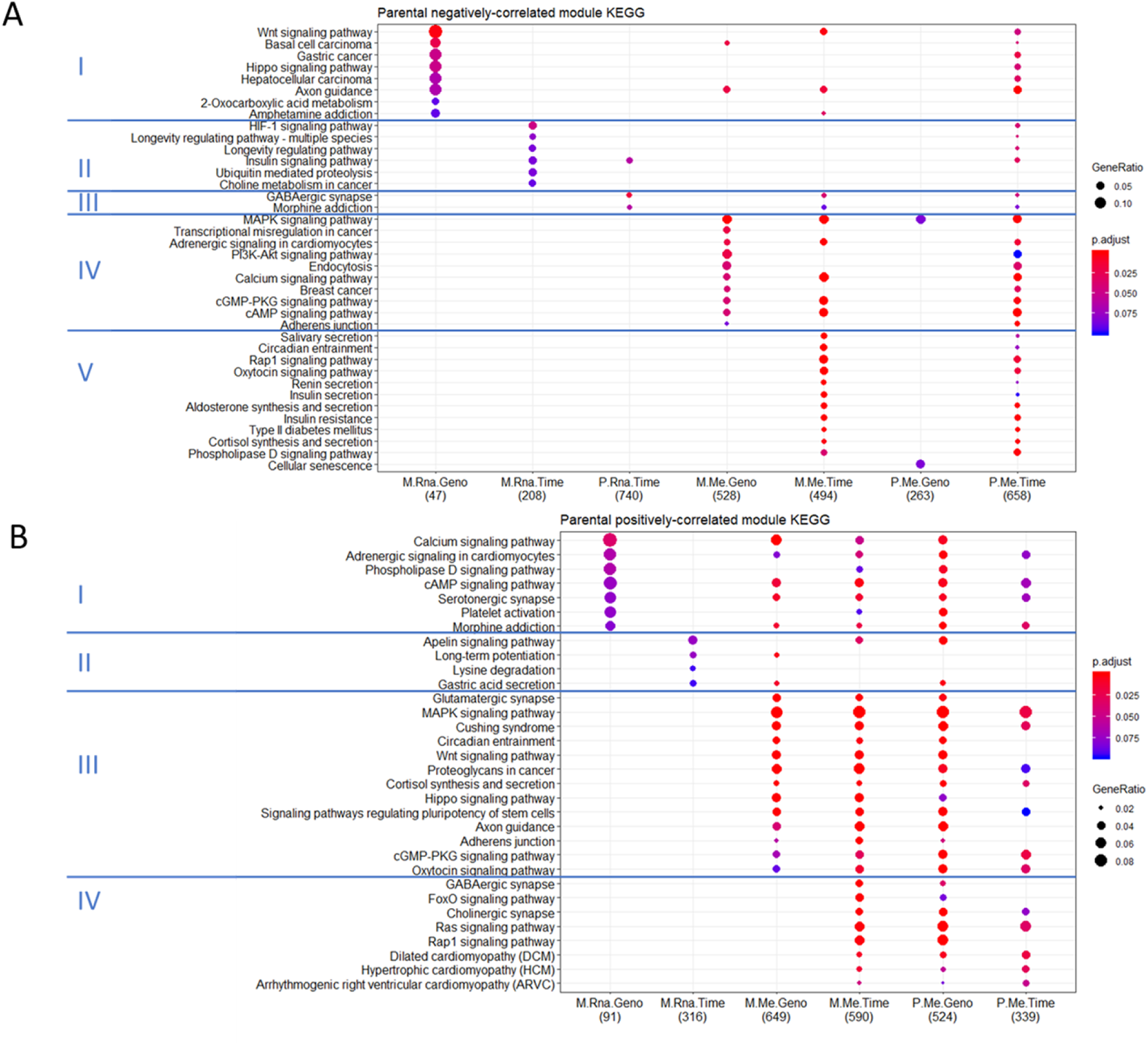
Cluster profiling of functional KEGG pathway enrichments for co-expression and co-methylation analyses. Comparison of KEGG profiles for all datasets using clusterProfiler on **A)** negatively-correlated and **B)** positively-correlated modules. Top 10 KEGG enrichment pathways with Benjamini-Hochberg adjusted p-values for each dataset are shown. M.Rna.Geno – maternal co-expression genotype-significant module genes. M.Rna.Time - maternal co-expression time-significant module genes. P.Rna.Time - paternal co-expression time-significant module genes. M.Me.Geno – maternal co-methylation genotype-significant module genes. M.Me.Time – maternal co-methylation time-significant module genes. P.Me.Geno – paternal co-methylation genotype-significant module genes. P.Me.Time - paternal co-methylation time-significant module genes. Full lists of GO terms and associated genes are in S17 Data (RNA) and S18 Data (methylation).

**Fig 8.**
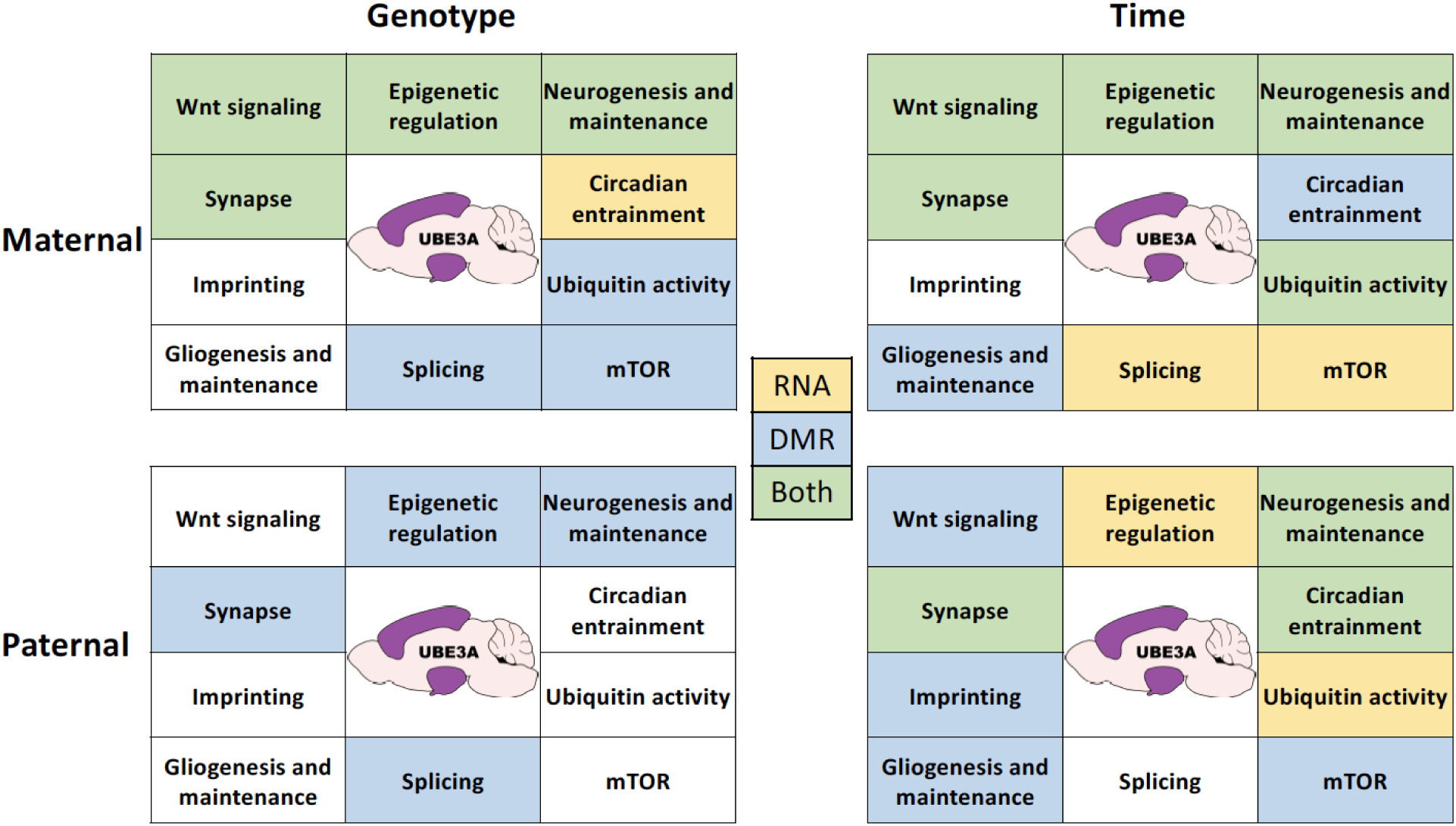
Network enrichment summary. General network categories enriched in WGCNA modules across time and genotype for maternal-derived and paternal-derived deletion samples. Enrichment is denoted for each data set for co-expression modules (yellow), co-methylation modules (blue), or both data sets (green).

## Discussion

Together, our data reveals the genome- and epigenome-wide influence of UBE3A on co-regulated genetic networks in a novel rat model with a complete CRISPR/Cas9 knockout of *Ube3a*. We demonstrated that the *Ube3a*-deletion rat model displays the expected transcript levels based on parental imprinting effects in two brain regions. Current AS mouse models contain either a single exon mutation of the maternal *Ube3a* (30), which is not a complete knockout of *Ube3a* similar to the majority of human AS patients, or a large deletion containing part of *Ube3a* extending to *Gabrb3* (39), which confounds the effect of *Ube3a* itself. Moreover, functional AS-relevant phenotypes have been largely dependent on laboratory and background strain of the model (40,41).Given the above considerations, our CRISPR/Cas9 *Ube3a* knockout rat model provides an improved model to study the effects of complete loss of maternal UBE3A as well as the effects of imprinting at this region by reciprocal parental deletions.

Single-gene approaches to analyze the effect of *Ube3a* deletion, such as DGE and DMR analysis, identified specific genes involved in neuronal processes that are affected transcriptionally and epigenetically by loss of both *Ube3a* sense and antisense transcripts. To better understand the co-expression and co-methylation gene network complexities related to *Ube3a*’s parental imprinting and developmental age effects on gene pathways, we also used the systems biology approach of WGCNA. Using WGCNA and associated GO term clustering approaches, we demonstrate that maternal *Ube3a* deletion predominately results in the transcriptional dysregulation of Wnt signaling and other developmental pathways. These results are consistent with the expected maternally inherited phenotypes associated with AS, as well as the brain anatomical differences that were specific to the maternal *Ube3a* deletion offspring in this same AS rat model (37). In this related study, maternal *Ube3a* deletion rats showed significantly smaller cortical regions and significantly larger hypothalamic brain regions, which is consistent with the dysregulated gene pathways involved in developmental and signal transduction pathways observed in our analyses of these two brain regions with maternal *Ube3a* inheritance.

In contrast to the dysregulated gene modules identified by RNA-seq, which were predominantly associated with maternal *Ube3a* deletion, those identified by WGBS methylation analyses were altered mostly equivalently by either maternal or paternal *Ube3a* deletion. These results are consistent with a model in which the maternally expressed protein product UBE3A is upstream of the developmental transcriptional dysregulation in AS, but the noncoding *Ube3a-Ats* regulates reciprocal changes to DNA methylation patterns on both maternal and paternal alleles.

We identified biologically relevant gene networks associated with differential developmental age and genotype in both parentally-derived *Ube3a* deletions. Together, these modules shared enrichments for pathways involved in Wnt signaling, epigenetic regulation, synaptic processes, imprinting, neuro and gliogenesis, circadian entrainment, ubiquitin activity, mTOR, and splicing (**Fig 6**). Enriched pathways in paternal samples were also seen in the maternal samples including Wnt signaling, ubiquitin, mTOR, neurogenesis, synapse, splicing, and epigenetic regulation. Additionally, direct comparison of the functional profiles revealed distinct patterning of network-wide parent-of-origin effects across genotype and developmental time and between the transcriptome and methylome. This approach has revealed that *Ube3a* deletion impacts specific neuronal networks and cellular processes at both the transcriptional and epigenetic level and that the paternally expressed *Ube3a-Ats* transcript has an epigenetic, gene-regulatory function in the developing brain that extends beyond its silencing effect on *Ube3a* sense expression.

Our systems biology approach has identified broad gene networks affected by UBE3A; however, the biological significance of the individual genes identified should be validated by more focused, functional studies. How the pathways identified here might inform about the observed behavioral phenotypes will be important to explore in future studies in order to identify novel treatment strategies to ameliorate disease phenotypes. Another limitation of this this study is the lack of separation of specific cellular subpopulations of the brain regions. The entirety of the hypothalamus was utilized and a whole hemisphere of cortex was used without sorting of neuron and glial populations. Since *Ube3a* is not imprinted in glia, the loss of this low level of paternal *Ube3a* expression in glia may have had an impact on our analyses. In future studies, single-cell sequencing approaches to capture expression and methylation differences within distinct cellular subpopulations could provide further insight into the genome-wide impacts of UBE3A loss in AS models.

The mechanism by which *Ube3a* is silenced by *Ube3a-Ats* is still not completely understood. *Ube3a-Ats* is at the end of a larger transcript where several additional noncoding RNAs with functions lacking in Prader-Willi syndrome are processed, particularly the *Snord116* snoRNA and its host gene *116HG* (*SNHG14*). Interestingly, both maternal and paternal *Ube3a* deletion resulted in dysregulated gene pathways that overlapped with those observed with paternal *Snord116* deletion in mouse cortex, including circadian entrainment, mTOR, and imprinting (5,6). Together with the results of this study, the evidence suggests that the paternally expressed *Ube3a-Ats* transcript may have similar but non-overlapping functions with *SNHG14* and that both maternal and paternal alleles may co-operate in the regulation of circadian rhythms of mTOR-mediated metabolism. These results also fit the recent hypothesized co-evolution of imprinted genes and rapid eye movement (REM) sleep patterns in mammals (42,43). Overall, these results suggest a possible cross-talk between maternal and paternal alleles of the AS/PWS loci in regulating DNA methylation changes implicated in the entrainment of circadian rhythmicity in neurons (44).

## Materials and Methods

### Rat breeding and sample collection

Cas9 and gRNA complexes were injected into fertilized Sprague Dawley rat embryos and inserted into a surrogate. Founders were used to produce F1 progeny and animals were screened for UBE3A expression. Samples were bred in reciprocal crosses from deletion carrier parents to produce maternal and paternal-derived deletion pups. While no sex differences are observed in Angelman syndrome, we chose to investigate females only to avoid the confounding variable of sex in genomic investigations. Female pups were harvested for tissue at P2 and P9 from at least 3 litters. Hypothalamus and cortex were collected and flash frozen and tail snips were collected for genotyping. Tail snips were prepared using the REDExtract-N-Amp Tissue PCR Kit (Sigma-Aldrich XNAT) and were screened for *Ube3a* deletion using PCR primers spanning the expected deletion site. Brain tissue samples were used for DNA and RNA extraction using the ZR-Duet DNA/RNA MiniPrep Plus kit (Zymo Genesee 11-385P). All procedures were approved by the UC Davis IACUC committee, protocol # 20035.

### Library preparation and sequencing

RNA from P2 and P9 hypothalamus and cortex was prepared for RNA-seq library using the NEBNext rRNA Depletion kit (NEB #E6310) and NEBNext Ultra II Directional RNA Library Prep Kit for Illumina (NEB #E7760S). Libraries were assessed for quality control on the Agilent Bioanalyzer 2100. Maternal and paternal libraries were prepared separately then pooled for multiplex sequencing and run 12 samples per lane PE 150 on the HiSeq4000.

DNA from P2 and P9 hypothalamus was used for WGBS. DNA was bisulfite converted using the EZ DNA Methylation-Lightning Kit (Zymo D5031) and prepared for WGBS libraries using the TruSeq DNA Methylation Kit (Illumina EGMK91324). Libraries were assessed for quality control on the Agilent Bioanalyzer 2100. Maternal and paternal samples were were prepared separately then pooled for multiplex sequencing and run 12 samples per lane PE 150 on the HiSeqX.

### RNA-seq data processing and DGE

Raw fastq files were processed and trimmed using HTStream (https://github.com/ibest/HTStream) then used as input for stranded PE alignment usingSTAR (https://github.com/alexdobin/STAR) (45) to output gene count files for each sample. Raw read counts were used for DGE using edgeR (46,47). Briefly, reads were filtered for one count per million in at least 6 libraries, libraries were normalized using the weighted trimmed mean of M-values (TMM), and fit to a single, combined model and extracted pairwise coefficients. DE genes from comparisons were used for GO enrichment analysis using Enrichr (48,49).

### WGBS data processing and DMR analysis

Raw fastq files were processed and trimmed of adapters and methylation bias (m-bias) using the CpG_Me repository (https://github.com/ben-laufer/CpG_Me) (50,51) to generate quality control reports and Bismark cytosine reports for DMR analysis. DMRs and other methylation analyses were performed using the DMRichR package and executable (https://github.com/ben-laufer/DMRichR) (50,52,53). Briefly, this DMR analysis approach utilizes statistical inference (smoothing) to weight CpG sites based on their coverage. CpGs in physical proximity that also show similar methylation values were then grouped into testable background regions that also showed a difference between groups, and a region statistic for each was estimated. Statistical significance of each testable background region was determined by permutation testing of a pooled null distribution. Annotated DMRs were used for GO enrichment analysis using Enrichr.

### Network analysis

Network analysis and module detection was performed using WGCNA (Zhang and Horvath 2005; Langfelder and Horvath 2008). For co-expression, RPKM was used as input, whereas for co-methylation, background smoothed methylation values output from DMRichR were used. Significant trait modules were subjected to GO enrichment analysis using Enrichr, where co-methylated modules were first annotated using ChIPseeker (54). KEGG enrichment profiling was done using the R package clusterProfiler (55). Ensembl gene ID’s for each data set were obtained using rat annotation and KEGG pathway enrichment was tested using p-value cutoff of 0.1 and Benjamini-Hochberg q-value cutoff of 0.5. Top 10 pathways for each group were output.

## Data Availability

Sequencing data are available in GEO, accession number pending. All the numerical data for the figures and statistics are provided in Supporting Information.

## Author contributions

Data Curation and Formal Analysis by SJL and BIL; Conceptualization, Supervision, and Funding Acquisition by JLS, DJS, and JML; Investigation and Methodology by SJL, BIL, UB, and ELB; Project administration by SJL, DJS, and JML; Validation, Visualization and Writing - original draft by SJL; Writing – review and editing by BIL, JLS, DJS, and JML.

## Supporting Information

**S1 Figure. Design for CRISPR-mediated UBE3A deletion rat model**. Location of gRNA’s for targeted deletion of UBE3A using 2 upstream 5’ gRNAs and 2 downstream 3’ gRNAs.

**S2 Table. Pairwise differentially expressed genes comparison groups and differentially expressed transcript numbers (p<0.05)**

**S3 Figure. Differentially expressed genes are largely unique between pairwise groups**

**S4 Data. Differentially expressed genes lists for all pairwise comparisons**.

**S5 Table. WGBS comparison groups and differentially methylated region numbers**

**S6 Figure. Differentially methylated region detection and clustering**. Clustering of DMRs identified by DMRichR for A) maternal time, B) maternal genotype, C) paternal time, and D) paternal genotype. DMRs along the y-axis are clustered against tested sample dendrograms on the x-axis. Samples were weighted for the trait of interest using the other as a covariate.

**S7 Figure. DMR Manhattan-plot distribution**. DMR distribution across the genome. Chromosome location is displayed across the x-axis and significance by permutations test on the y-axis. Heatmap along the x-axis displays the number of CpGs in each DMR.

**S8 Data. List of genes and statistical information on pairwise differentially methylated regions**

**S9 Figure. Sample cluster dendrograms for WGCNA**. Sample clustering for RPKM in hypothalamus for A) maternal samples and B) paternal samples, in cortex for C) maternal samples and D) paternal samples, while samples clustering for background smoothed methylation values in hypothalamus for E) maternal samples and F) paternal samples. Control sets were conducted using the wildtype samples for both maternal and paternal samples using RPKM in G) hypothalamus and H) cortex, and I) background smoothed methylation in hypothalamus.

**S10 Figure. Cortex WGCNA significant module trait heatmap and GO terms.** Co-expression modules detected from network analysis of RPKM in A) maternal and B) paternal deletion cortex. Modules were grouped by co-expression and tested for correlation with the traits of genotype or time. Modules of interest are correlated (red) or anticorrelated (blue) with each trait (p-value < 0.05, highlighted purple and yellow boxes). Highlighted co-expression module gene lists were used for GO enrichment with Enrichr using GO Biological Process 2018, GO Cellular Component 2018, GO Molecular Function 2018, KEGG 2016, and MGI Mammalian Phenotype Level 4 categories. Top GO categories are displayed as combined score, calculated as - log10(p-value)*z-score.

**S11 Figure. WGCNA and GO of control wild-type litter effects.** Co-expression modules detected from network analysis of RPKM between maternal and paternal wild-type control samples for hypothalamus and cortex. Modules were grouped by co-expression and tested for correlation with the traits of parent (litter-mates of maternal vs. paternal deletion) or time. Modules of interest are correlated (red) or anticorrelated (blue) with p-value < 0.05 (highlighted purple and yellow boxes). Highlighted co-expression module gene lists were used for GO enrichment with Enrichr using GO Biological Process 2018, GO Cellular Component 2018, GO Molecular Function 2018, KEGG 2016, and MGI Mammalian Phenotype Level 4 categories. Top GO categories are displayed as combined score, calculated as -log10(p-value)*z-score.

**S12 Figure. Pair-wise hypothalamic differential gene expression gene ontologies by time or parental origin of *Ube3a* genotype**. Enrichr gene ontology enrichment for GO Biological Process 2018, GO Cellular Component 2018, GO Molecular Function 2018, KEGG 2016, and MGI Mammalian Phenotype Level 4 categories. Ontologies for gene lists obtained from DGE lists for **A)** maternal time comparisons, **B)** maternal genotype comparisons, **C)** paternal time comparisons, and **D)** paternal genotype comparisons. Combined score is calculated as -log10(p-value)*z-score.

**S13 Figure. Pair-wise hypothalamic differential methylation gene ontology enrichment by time or parental origin of *Ube3a* genotype**. DMR-associated genes from each data set were used for gene ontology enrichment with Enrichr using GO Biological Process 2018, GO Cellular Component 2018, GO Molecular Function 2018, KEGG 2016, and MGI Mammalian Phenotype Level 4 categories. Top ontology terms are displayed for differential methylation with **A)** maternal time, **B)** maternal genotype, **C)** paternal time, and **D)** paternal genotype. Combined score is calculated as -log10(p-value)*z-score.

**S14 Data. Significant WGCNA module gene lists for all groups.**

**S15 Cluster profiling of functional GO enrichments for negatively-correlated co-expression and co-methylation analyses**. Comparison of negative-correlation modules GO profiles for all datasets using clusterProfiler. GO enrichment categories used were Biologic Process, Cellular Function, and Molecular Function with Benjamini-Hochberg adjusted p-values. Top 10 categories for each dataset are shown. M.RNA.Geno – maternal co-expression genotype-significant module genes. M.RNA.Time - maternal co-expression time-significant module genes. P.RNA.Time - paternal co-expression time-significant module genes. M.DMR.Geno – maternal co-methylation genotype-significant module genes. M.DMR.Time - maternal co-methylation time-significant module genes. P.DMR.Geno – paternal co-methylation genotype-significant module genes. P.DMR.Time - paternal co-methylation time-significant module genes.

**S16 Figure. Cluster profiling of functional GO enrichments for positively-correlated co-expression and co-methylation analyses.** Comparison of positive-correlation modules GO profiles for all datasets using clusterProfiler. GO enrichment categories used were Biologic Process, Cellular Function, and Molecular Function with Benjamini-Hochberg adjusted p-values. Top 10 categories for each dataset are shown. M.RNA.Geno – maternal co-expression genotype-significant module genes. M.RNA.Time - maternal co-expression time-significant module genes. P.RNA.Time - paternal co-expression time-significant module genes. M.DMR.Geno – maternal co-methylation genotype-significant module genes. M.DMR.Time - maternal co-methylation time-significant module genes. P.DMR.Geno – paternal co-methylation genotype-significant module genes. P.DMR.Time - paternal co-methylation time-significant module genes.

**S17 Data. Significant co-expression module genes GO for all groups**.

**S18 Data. Significant co-methylation module genes GO for all groups**.

